# Integrative deep learning analysis improves colon adenocarcinoma patient stratification at risk for mortality

**DOI:** 10.1101/2022.06.13.495227

**Authors:** Jie Zhou, Ali Foroughi pour, Hany Deirawan, Fayez Daaboul, Thazin Aung, Rafic Beydoun, Fahad Shabbir Ahmed, Jeffrey H. Chuang

## Abstract

Colorectal cancers are the fourth most commonly diagnosed cancer and the second leading cancer in number of deaths. Many clinical variables, pathological features, and genomic signatures are associated with patient risk, but reliable patient stratification in the clinic remains a challenging task. Here we assess how image, clinical, and genomic features can be combined to predict risk. We first observe that deep learning models based only on whole slide images (WSIs) from The Cancer Genome Atlas accurately separate high risk (OS<3years, N=38) from low risk (OS>5years, N=25) patients (AUC=0.81±0.08, 5year survival p-value=2.13e-25, 5year relative risk=5.09±0.05) though such models are less effective at predicting OS for moderate risk (3years<OS<5years, N=45) patients (5year survival p-value=0.5, 5year relative risk=1.32±0.09). However, we find that novel integrative models combining whole slide images, clinical variables, and mutation signatures can improve patient stratification for moderate risk patients (5year survival p-value=6.69e-30, 5year relative risk=5.32±0.07). Our integrative model combining image and clinical variables is also effective on an independent pathology dataset generated by our team (3year survival p-value=1.14e-09, 5year survival p-value=2.15e-05, 3year relative risk=3.25±0.06, 5year relative-risk=3.07±0.08). The integrative model substantially outperforms models using only images or only clinical variables, indicating beneficial cross-talk between the data types. Pathologist review of image-based heatmaps suggests that nuclear shape, nuclear size pleomorphism, intense cellularity, and abnormal structures are associated with high risk, while low risk regions tend to have more regular and small cells. The improved stratification of colorectal cancer patients from our computational methods can be beneficial for preemptive development of management and treatment plans for individual patients, as well as for informed enrollment of patients in clinical trials.

## Introduction

Stratification of colon adenocarcinoma patients is based on standards established by the American Joint Committee on Cancer (AJCC) and Union for International Cancer Control (UICC)^1^ and remains a challenging clinical decision. Colon adenocarcinoma has an overall all-stages SEER 5-year survival of 63% ^2,3^, and risk assessments impact decisions such as whether a patient receives additional chemotherapy or is inducted into a clinical trial. Tumor infiltrating lymphocytes (TIL) quantifications have been shown to be informative in recent years ^4-6^. Nevertheless, improvements in biomarkers remain critical, either through incorporating additional biomarkers or better use of currently identified markers^7-9^, particularly for patients without a clear indication of high/low risk^10,11^. Clinical assessment of these patients can be difficult, hampering decisions about additional treatment, uptake of patients into clinical-trials, and proactive disease surveillance^12,13^. Therefore, automated computational models on patient data, including histopathology images, can address important needs in assessment and reproducibility of cancer management decisions.

Deep learning models have achieved high accuracy for detecting tumor regions^14^ and identifying cancer subtypes^15^ from hematoxylin and eosin (H&E)-stained whole slide images. Such models have also been able to predict several clinically relevant genetic features, such as microsatellite instability (MSI) ^16^ and mutation status of key genes^17-19^ with moderate accuracy. Deep learning models using WSIs have been studied to stratify patients based on survival risk^20^. However, these models have room for improvement as they have tended to only utilize image features, have moderate AUCs, and rely on large datasets for model training, including in recent studies of colorectal cancer ^21,22^.

We hypothesize that integrating H&E image data with other data modalities can improve risk stratification since clinical variables, mutation status, and gene expression profiles have individually been shown to be informative ^23^. To address this question, we develop and evaluate integrative deep learning models that combine morphological features from H&E WSIs, clinical variables, MSI-status, and mutation status of key genes^24-31^. While prior studies have combined patient-level image features from WSIs with patient-level clinical variables, to the best of our knowledge, our work is the first to train at the tile level with patient-level information based on context-aware learning^32^, which we find improves performance.

We show that integrative analysis improves patient stratification and enables training of reliable models using smaller sample sizes, which we demonstrate using TCGA-COAD (The Cancer Genome Atlas - Colonic Adenocarcinomas)^33^ and an independently generated dataset. Our integrative model demonstrates superior performance to models using only one data type and is more robust to staining differences than a model using only WSIs. By combining local image features with patient level data our model outputs interpretable heatmaps, which are informative of morphologies of high risk. These results demonstrate how integrative computational analysis of colorectal adenocarcinomas can improve prediction of outcomes.

## Materials and Methods

### Data and study design

#### TCGA-COAD cohort

336 Formalin-Fixed Paraffin-Embedded (FFPE) hematoxylin and eosin (H&E) stained TCGA-COAD WSIs were downloaded from the GDC (Genomic Data Commons) data portal. The following clinical variables were downloaded from the cBioPortal webpage^34,35^: patient age at diagnosis, gender, tumor (T) stage, nodes (N) stage, and metastasis (M) stage. Mutation statuses of 207 genes were downloaded from the cBioPortal webpage (see Table S1 for the full gene list). Patients were grouped by their overall survival (OS): low-risk (LR, OS>5 years, N=25), moderate-risk (MR, 3<OS<5 years, N=45), high-risk (HR, OS<3 years, N=38), and loss to follow-up (time to last follow up <3 years, patient status: alive, N=228 censored patients). These 228 censored patients were used for the training of the computational tumor detector, since they comprised a set independent from the survival studies.

#### Wayne State University Validation Cohort

123 patients’ H&E stained FFPE samples and corresponding clinical data were collected from Wayne State University (WSU). The clinical data include patient age at diagnosis, gender, T stage, N stage and M stage. Patients were grouped as HR(N=17), LR(N=97) and MR(N=9). There was no loss to follow-up in this cohort.

#### Combined multi-center cohort

A multi-center cohort (N=115) was obtained by combining the WSU (N=43) validation cohort and TCGA-COAD (N=72). TCGA-COAD slides obtained from Indivumed (N=36) were removed (see Model training and assessment section). Patients were grouped as HR(N=39), LR(N=42) and MR(N=34).

### Data preparation

#### Tumor annotation

Tumor regions of WSIs were annotated by expert pathologists using the Aperio ImageScope software version 12.4.3^36^. Our pathologists only annotated highly pure tumor regions. Tumor areas were exported from the Aperio software in The Extensible Markup Language (XML) format, with X and Y coordinates corresponding to the annotated tumor regions. Tumor masks were generated for each slide image by connecting the coordinates, dilating, and eroding the areas using the OpenCV package in python^37^.

#### Image pre-processing

H&E stained FFPE WSIs were acquired in SVS format. All images were downsampled to 20× magnification, corresponding to a resolution of 0.5 μm/pixel. Each WSI was manually reviewed and the tumor area was annotated by expert pathologists. Regions with excess background or containing no tissue were removed as previously described ^38^. Image slides were tiled into non-overlapping patches of 512 × 512 pixels. Tiles with >50% overlap with tumor masks were labeled as tumor tiles. The remaining tiles were labeled non-tumor.

#### Clinical data pre-processing

Five clinical variables related to patient outcomes were selected by the pathology team: age at diagnosis, gender, and TNM (Tumor, Nodes, and Metastasis) staging of colonic adenocarcinomas:tumor (T) stage, nodes (N) stage, and metastases (M) stage based on the college of American pathology (CAP) protocol for Colon and Rectum, Resection 2021 (v4.2.0.0)). Age was encoded numerically, and other variables were encoded as integers.

#### Molecular data pre-processing

207 genes from 11 canonical cancer pathways ^24-31^ and the top 11 most commonly mutated genes in TCGA-COAD were selected. A 10% threshold was used to filter out genes that are not frequently mutated in TCGA-COAD patients, resulting in a total of 26 genes (see Table S1). Microsatellite instability (MSI) status was also considered due to its impact in colon cancers ^39^. Table 1 depicts detailed patient characteristics for the data sets.

**Table 1.** Survivorship of High/Low risk patients in TCGA test set.

| High/Low | 5-year LR | 5-year HR | 5-year *p*-value |
| --- | --- | --- | --- |
| Image-only | 0.615 | 0.296 | 2.13E-25 |
| Image & clinical | 0.645 | 0.189 | 1.92E-23 |
| Image & mutation | 0.524 | 0.283 | 1.37E-21 |
| Image, clinical & mutation | 0.658 | 0.324 | 1.20E-19 |
| Clinical-only | 0.548 | 0.271 | 1.03E-24 |
| Mutation-only | 0.554 | 0.313 | 1.58E-20 |
| Clinical & mutation | 0.552 | 0.266 | 5.42E-25 |

### Model training and assessment

#### Train-test splits and cross-validation

We used Monte Carlo cross-validation to assess the model performance. We randomly split our cohort into paired training (70%) and testing (30%) sets to generate 100 training/testing set pairs. The predictive accuracy was assessed in each split. The results were then averaged over the splits.

#### Network architecture

InceptionV3^40^ features pre-trained on Image-Net^41^ were fed to a two-layer multi-layer perceptron (MLP) following the parameters of ^38^: The first layer has 1024 neurons followed by ReLU activation and drop-out. The second layer is the classification layer with softmax activation. Parameter initialization and batch size (=512) was set according to ^38^ L1-L2 regularization values and number training epoch were the two hyper-parameters optimized over subsets of data before the final cross validation step (number of epochs =10, L1-L2 regularization, regularization of 10e-4 for both L1 and L2 penalties. The model only using WSIs was used for hyper-parameter optimization.

For the computational tumor detector we used the same architecture from ^38^.

#### Feature construction

Mutation status was encoded as a binary variable. Age was encoded as a continuous variable. Other clinical variables were encoded as integers as well as one-hot-encoded variables. The deep learning survival model used the one-hot-encoding outputs. Random forest models trained on clinical and mutation status considered both encodings. Integer encoding resulted in higher AUCs and was used throughout. Clinical and/or mutation status variables were concatenated with the tile level Inception V3 features (see Figure 1).

**Fig. 1.**
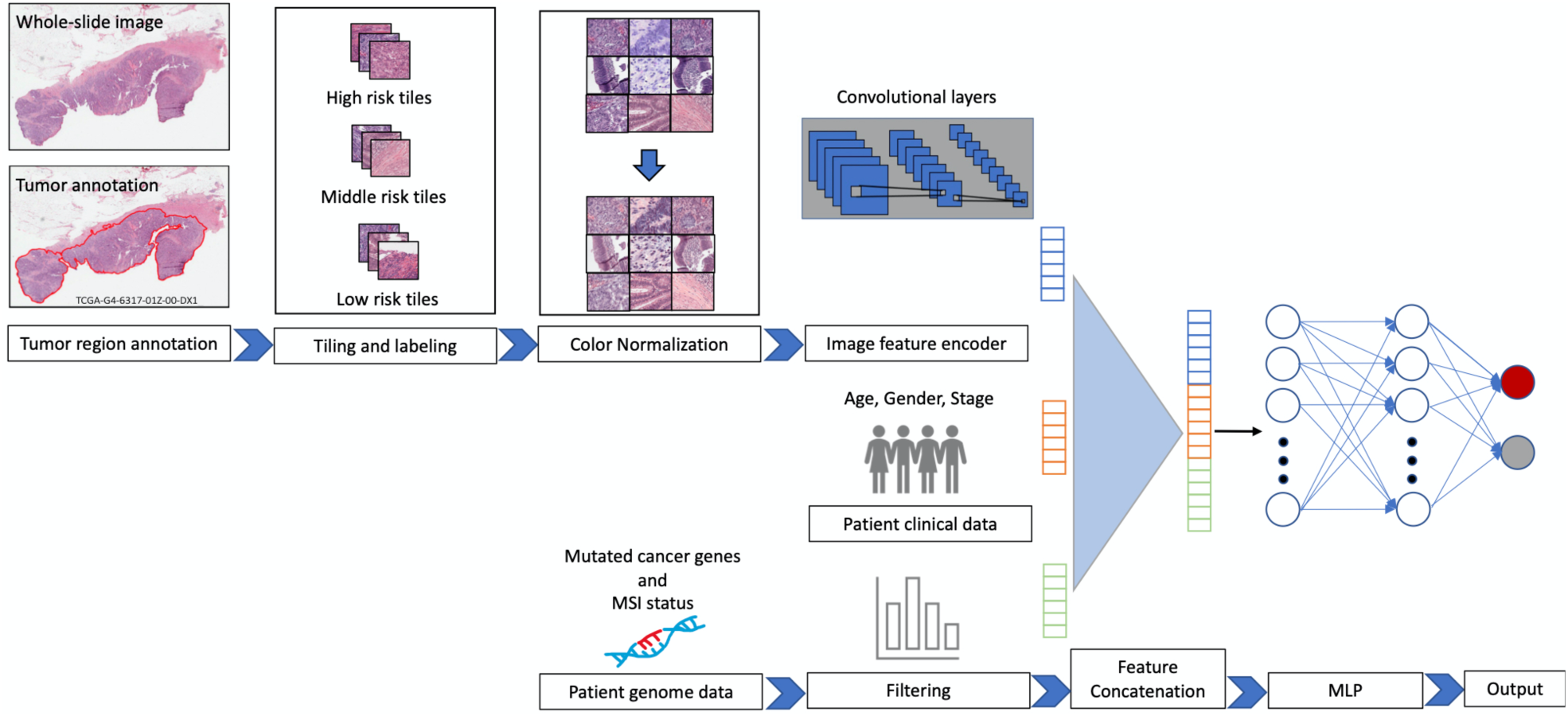
The integrative CNN model. Tumor regions of WSIs are annotated by expert pathologists. WSIs are tiled, and tiles overlapping with pathologist tumor annotations (>50% overlap) are used for survival analysis. Tiles are color normalized using the Macenko^42^ method and passed through an Inception V3 model pre-trained on Image-Net. Tile level CNN features are concatenated with patient level clinical variables and mutation status. These features are fed to a multi-layer perceptron to predict patient risk.

#### Deep learning model training

Deep learning models were trained to predict risk either from WSIs only, or as integrative models that combine WSIs and other data modalities. A key difference of our method compared to others is how we use local information in the training of the integrative models. In previous approaches^21^ tile-level image features are averaged within a patient to create patient-level image features. Patient-level image features are then used with patient level clinical variables to train the classification model. In our integrative models, however, each tile is concatenated with the patient clinical features, and training is done across all tiles. We did this because we found that using patient-level image features in the training yielded inferior performance (AUC=0.68±0.09 for deep learning Cox model and AUC=0.81±0.08 for image-only model, compare Figures 2b and Supplementary Figure 1i).

**Fig. 2.**
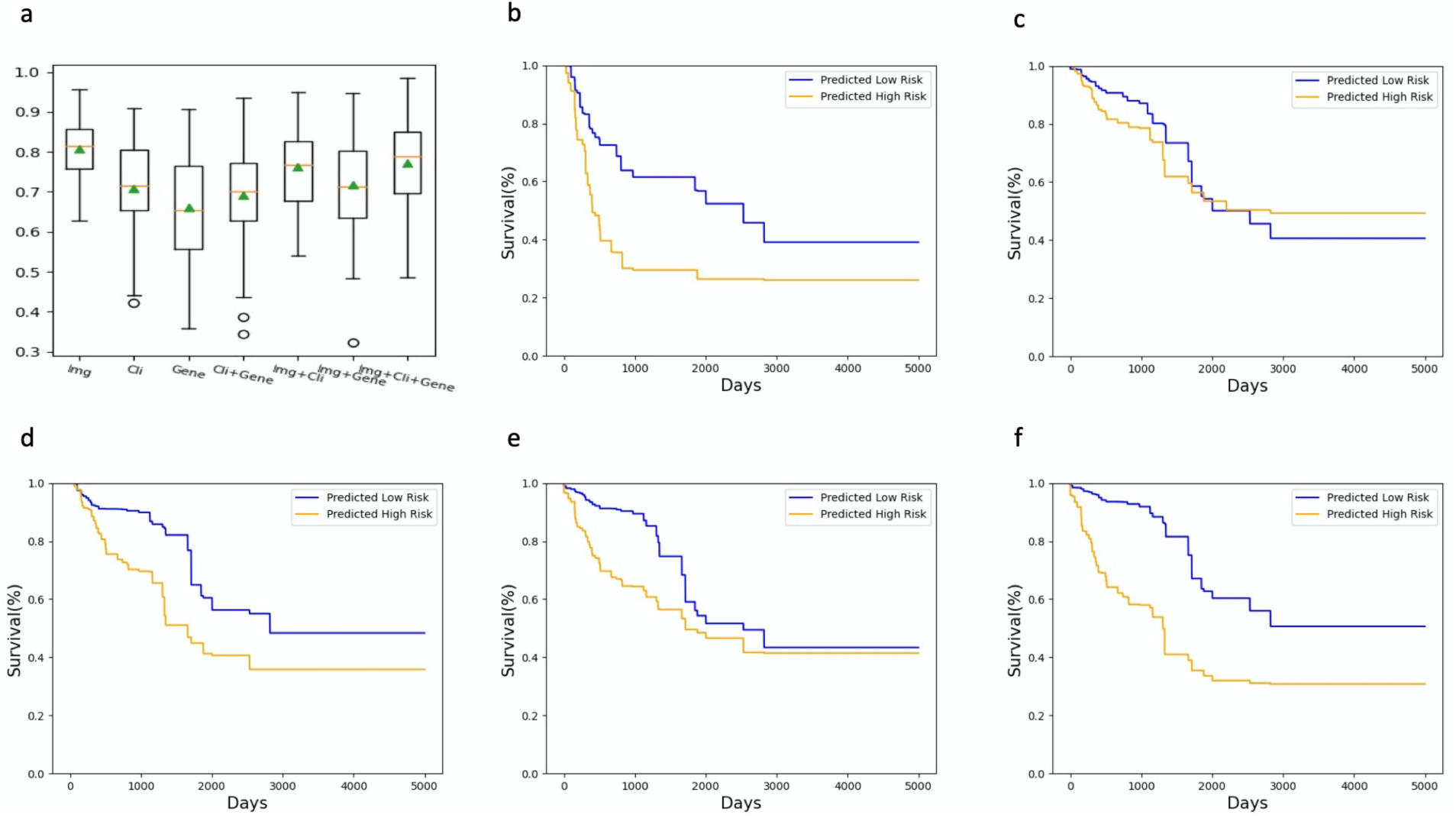
Integrative analysis improves stratification of moderate risk patients. (a) AUCs for prediction of High/Low risk class by various models. Kaplan-Meier curves of patients from the (b) High/Low and (c) High/Moderate/Low clinical groups, stratified by predicted risk class from the image-only model. Kaplan-Meier plots of High/Moderate/Low patients as stratified by the (d) image & clinical, (e) image & mutation, and (f) image & clinical & mutation models.

(1) Image-only model: for the image-based model, we utilized the Inception V3 architecture that was pretrained on the ImageNet database as described previously^38^. The cached 2048 global average pooling layer features of InceptionV3 were extracted and written to disk for downstream analysis. (2) Integrative models: we designed integrative prognostic models integrating WSIs with different combinations of data modalities. We concatenated tile-level InceptionV3 features with the feature vectors encoding clinical variables and/or mutation signature. The final feature vector was fed to the two-layer MLP. We under-sampled tumor tiles of the majority class to mitigate the effects of class imbalance. To address potential batch effects, we utilized the Macenko method^42^ to normalize the stain color across training and independent test data sets. (3) Deep learning Cox model: we trained a Cox proportional hazards model using patient-level image features extracted from Inception V3 transfer learning architecture. Specifically, slide-level image features were generated using the median value of all tumor tiles, and were fed into a Cox proportional hazards regression model implementation of the statsmodels (v0.13.2) package. The median hazard scores of patients in each class (HR or LR) of the train set were averaged, serving as the threshold on the hazard score for predicting class labels in the test set.

#### Random forest for survival analysis

We used random forests to train 3 separate models stratifying patients based on clinical variables, mutation status, and combined clinical and mutational signatures. Random forests were implemented using Sklearn version 1.02^43^ with default settings except the number of trees was set to 300. We tested several MLPs, but they performed inferior to the random forest model and were less stable. Therefore, the random forest model was used as the final classifier.

#### Model assessment

Model performance was evaluated on patients in the test set. Tile-level risk probabilities were averaged to construct patient level scores. A threshold of 0.5 was used to predict patients as High risk (HR) or Low risk (LR). No threshold optimization was performed. The Kaplan-Meier (KM) curves were plotted using the averaged survivorship at each time point in each cohort. 3year and 5year survivorships were used to assess model performance. In addition to KM plots, the mean and standard deviation of the area under the receiver operating characteristic (AUROC) on the test set was used to measure classifier performance in separating HR and LR patients.

#### Relative Risk Score

The mean and standard deviation of relative-risk at 3 year, 5 year, and median survival points were calculated to compare KM curves. For each test set relative-risk was calculated as follows, where SL and SH denote the survivorship of predicted LR and predicted HR patients, respectively:

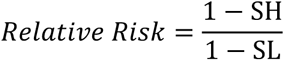

#### Feature importance assessment

We used SHAP (SHapley Additive exPlanations)^44 45,46^ to explain the predictions of our trained models. SHAP measures the impact of each feature value on the predictions of a machine learning model for a single input. The average SHAP impact across a dataset quantifies the overall variable importance for a fixed machine learning model. The KernelExplainer function of SHAP was used to measure importance of clinical variables and InceptionV3 features in the integrative deep learning model. 50 randomly selected tiles were used to estimate variable importance of the integrative model. The clinical-only model, being a random forest, uses the TreeExplainer function of SHAP to measure the importance of each clinical variable. Beeswarm plots depict the impact of top variables on each patient, and bar plots depict the average SHAP value magnitudes of top variables for each class. For each variable group total importance is defined as the sum of the importance of all variables in the group (e.g. all clinical variables or all InceptionV3 image features).

#### Image comparison across centers

The WSU validation cohort and TCGA-COAD cohort were compared to assess relative image quality and compatibility. We observed stronger differences between the WSU images and TCGA Indivumed slides than between the WSU images and TCGA-COAD slides from other TCGA centers (see Supplementary Figure 4). This difference was observable despite Macenko normalization. Removing Indivumed slides reduced stain differences and improved generalizability of our TCGA-models to WSU data. For this reason we removed Indivumed slides from the multicenter analysis as well. The outlier behavior of the TCGA Indivumed slides has been reported in prior studies of TCGA WSIs^47^ as well.

## Results

### 3.1 Images are informative of colon adenocarcinoma risk

We first investigated to what extent WSIs alone are predictive of patient risk in TCGA-COAD (Figure 1). We binned patients as high-risk (HR, OS<3years, N=38), moderate risk (MR, 3years<OS<5years, N=45), and low risk (LR, OS>5years, N=25) based on overall survival (see methods). We trained a convolutional neural network (CNN) to predict risk from WSIs, hereafter called the image-only model (see methods). The image-only model is able to distinguish HR and LR patients (AUC=0.81±0.08, see Figure 2a), and separates the survival curves (see Figure 2b and Table 1, p-value=4.06e-26, relative risk=2.09±0.09). The separation between survival curves decreases when MR(3<OS<5) patients are included in the test set (see Figure 2c). We found that binarization of the patients into HR and LR groups was important to the predictive success.

We then compared the image-only model to models based on clinical variables and/or mutation statuses (see methods). The image-only (AUC=0.81±0.08) model performance was superior to models using only clinical variables (clinical-only model, see Figure 2a and Supplementary Figure 1b, AUC=0.71±0.12) or only mutation status (mutation-only model, see Figure 2a and Supplementary Figure 1a, AUC=0.66±0.12), as well as to an integrative model combining clinical and mutation information (clinical & mutation model, see Figure 2a and Supplementary Figure 1c, AUC=0.69±0.11). These results indicate WSIs are a rich source of information for separating HR and LR patients. Similarly, the clinical-only, mutation-only, and clinical & mutation models were less effective than the image-only model in separating the survival curves when MR patients were included in the test set (see Supplementary Figure 1g-i).

### 3.2 Integrative analysis improves stratification of moderate risk patients

We next tested whether an integrative model combining WSIs, clinical variables, and mutation status, hereafter called the image & clinical & mutation model, would improve patient stratification. We found that the fully integrative model performs similarly to the image-only model in separating HR and LR patients (see Figure 2a), but performs superiorly when MR patients are included (compare Figures 2f and Supplementary Figure 1f). This finding is similar to the skin cancer results of ^48^, who reported that an integrative model had comparable performance to single-data-type models for distinguishing patients with strong survivorship differences, but that the integrative model provided additional benefit for low confidence cases.

We also investigated integrative models utilizing two data modalities (image & clinical and image & mutation models, see Figure 2d-e and Supplementary Figure 1c-e) for stratifying patients. The integrative models using only two data types were inferior to the image & clinical & mutation model, though the image & clinical model was superior to the image & mutation model. Both the image & clinical and image & mutation models outperform the clinical & mutation model (see Supplementary Figure 1d, 1e and 1i). As shown in Table 2, the image & clinical & mutation model provided stronger separation of survival curves than any other model at the 3-year and 5-year time points.

**Table 2.** Survivorship of H/M/L patients of TCGA test set.

| High/Moderate/Low | 3-year LR | 3-year HR | 5-year LR | 5-year HR | 3-year *p*-value | 5-year *p*-value |
| --- | --- | --- | --- | --- | --- | --- |
| Image-only | 0.836 | 0.786 | 0.586 | 0.564 | 1.96E-05 | 5.96E-01 |
| Image & clinical | 0.899 | 0.697 | 0.65 | 0.449 | 1.25E-20 | 1.33E-14 |
| Image & mutation | 0.894 | 0.644 | 0.591 | 0.496 | 7.70E-20 | 3.81E-12 |
| Image & clinical & mutation | 0.919 | 0.581 | 0.671 | 0.355 | 2.76E-30 | 6.69E-30 |
| Clinical-only | 0.687 | 0.663 | 0.418 | 0.275 | 3.72E-02 | 1.54E-21 |
| Mutation-only | 0.437 | 0.459 | 0.437 | 0.459 | 1.00E+00 | 1.00E+00 |
| Clinical & mutation | 0.822 | 0.768 | 0.63 | 0.443 | 6.19E-02 | 3.75E-20 |

### 3.3 Prediction heatmaps reveal the morphology associated with risk

We analyzed the prediction heatmaps of several representative TCGA-COAD slides to gain insight about the underlying morphologies that CNNs associate with risk (see Figure 3). These heatmaps were generated using the image & clinical & mutation model and show the risk probability for each tile as predicted by the CNN. Pathologist review suggests that nuclear shape, nuclear size pleomorphism, intense cellularity, and abnormal structures are indicative of high risk. Low risk tiles tend to have more regular and small cells ^49-52^.

**Fig. 3.**
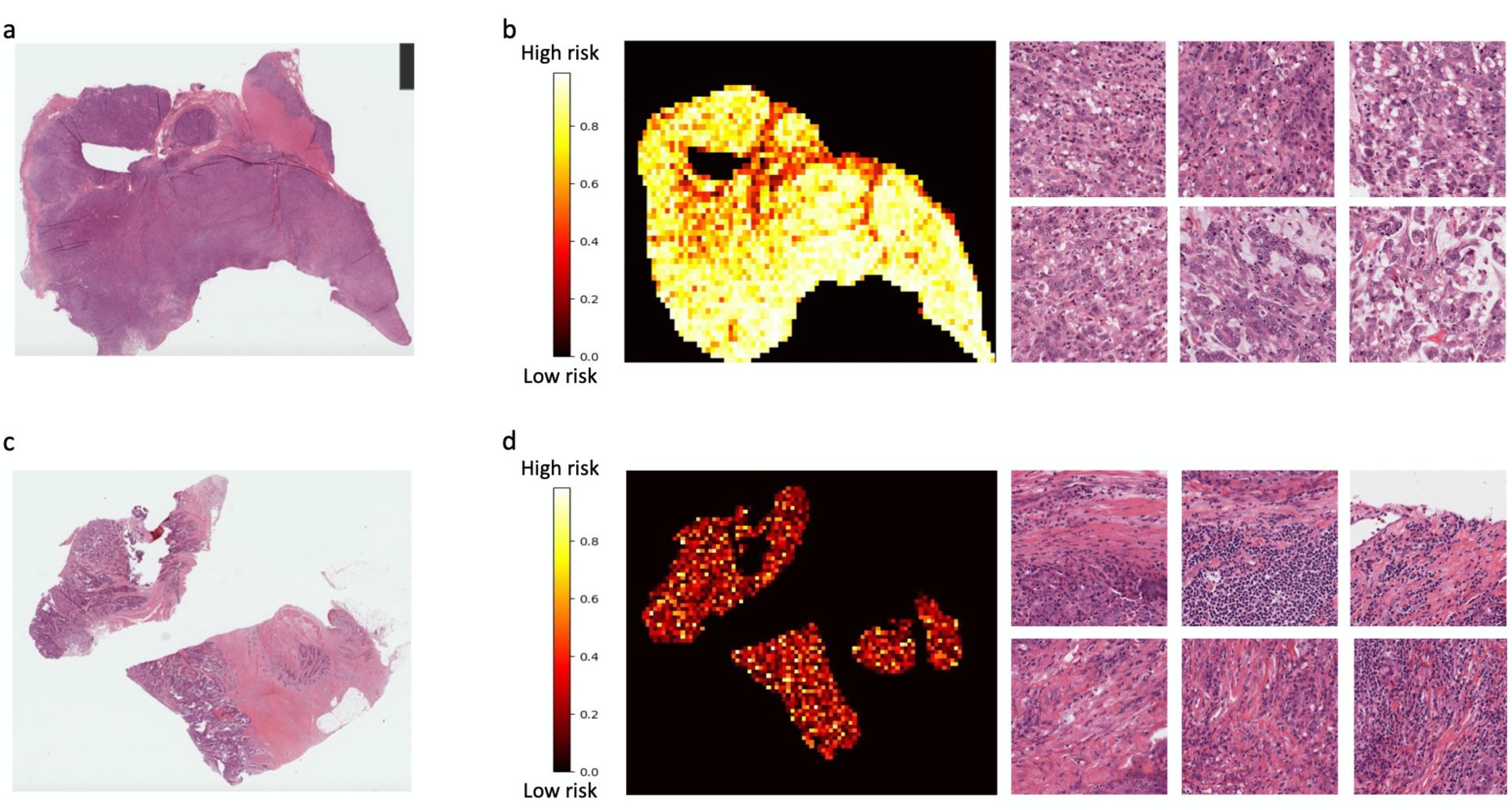
Representative H&E slides from TCGA test set and their predicted heatmaps. WSIs of (a) a high risk patient and (c) a low risk patient. The prediction heatmaps of (b-left) a high risk patient and (d-left) a low risk patient. Example tiles predicted as (b-right) high risk and (d-right) low risk from (a) high risk patient and (c) low risk patient, respectively.

### 3.4 Pure tumor regions are more informative of risk

Accurate identification of tumor regions within a WSI is a key preliminary step affecting risk classification. To test whether pathologist annotation of tumor regions can be replaced with a computational method, we used pathologist annotations of 228 independent WSIs (see Methods) to build a computational tumor detector. This detector showed high accuracy (Figure 4a, AUC >92%). Some other works have reported higher AUCs for computationally identifying tumor regions^53^, though this is likely due to variations in pathologist annotation methods. For example, some of our “false positives” are due to the fact that only some regions in a slide were selected for annotation (see Supplementary Figure 3). Other discrepancies between our computational predictions and pathologist annotations appear to be related to pathologists’ implicit thresholds for tumor annotation. Manual inspection of several “false positive” computational predictions indicate they do contain tumor cells but at lower purity than pathologist annotated regions.

**Fig. 4.**
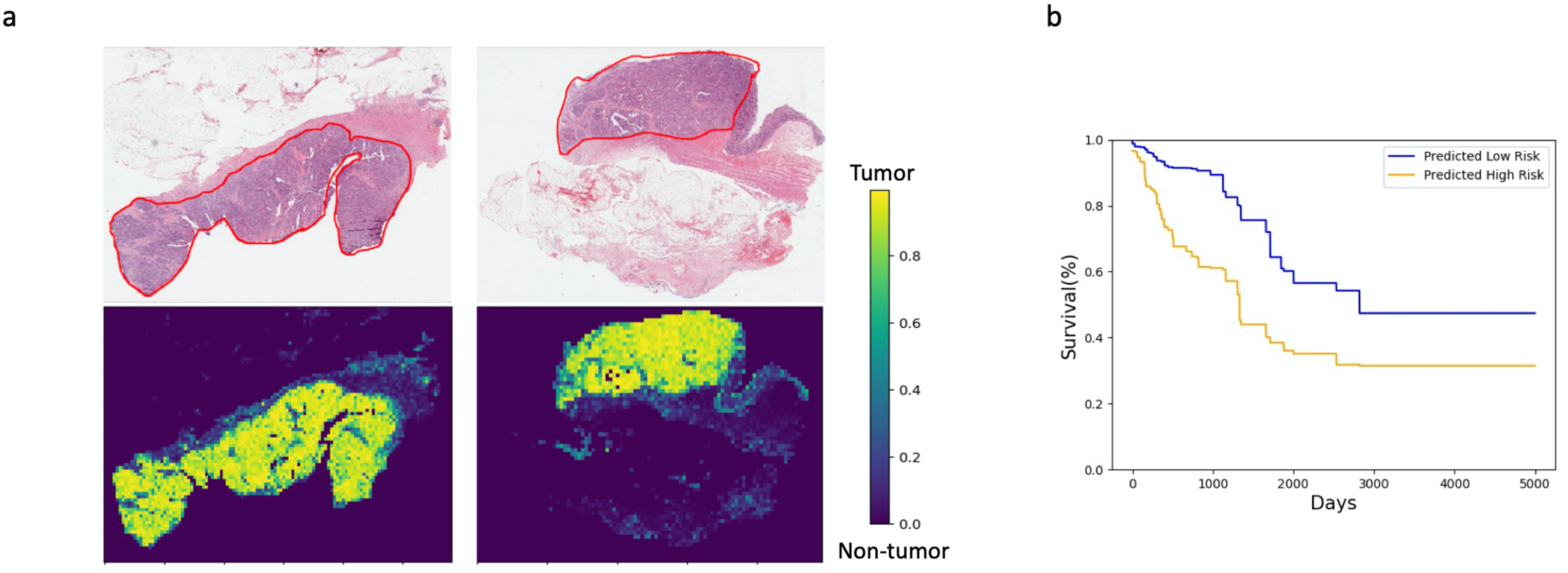
Accurate tumor detection improves survival prediction. (a) ground truth annotations of tumor regions from pathologists, circled with redlines (top) and tumor prediction heatmaps (bottom). (b) Kaplan-Meier curve of predicted high and low risk patients on the full set of High/Moderate/Low risk patients, determined by applying the image, clinical & mutation model to predicted tumor regions.

The Kaplan-Meier curve generated using our computational tumor detector as input into the image & clinical & mutation model to the H/M/L dataset is shown in Figure 4b. As in Figure 2f, there is a clear separation between the high and low risk curves. However, the separation is lower using the computational tumor detector than using pathologist annotations. The 3year relative risks when using pathologist annotated and deep learning-predicted tumor regions are 3.08 and 2.63, respectively (p-value < 0.05). These results suggest that the integrative model is more effective using only pure tumor regions as input, while computational tumor predictions tend to include low-purity regions that reduce performance.

### 3.5 Validation of TCGA models on Wayne State hospital data

As an additional verification, we validated our TCGA-trained models on an independent dataset from Wayne State University (WSU). We collected and annotated tumor regions (N=123, see materials and methods), and stratified patients as HR (N=17), LR(N=97), or MR(N=9) similar to the TCGA-COAD cohort. For analyses involving the Wayne State cohort, we did not include mutation data as it was not available.

We first considered a test set that included all HR, LR and MR cases together. The image-only model was unable to stratify high and low risk patients for this test set (3 year survival, image-only p-value=5.0e-01, Figure 5a). The clinical-only model provided a statistically significant but modest stratification (clinical-only p-value=3.3e-03, Figure 5b). However, the image & clinical model provided superior separation of the patient cohort (3 year survival, p-value=1.14e-09, see Figure 5c and Table 3), consistent with expectations from the intra-TCGA analysis.

**Table 3.** Survivorship of H/L patients of Wayne State data stratified by the TCGA-trained models.

| High/Low | 5-year LR | 5-year HR | 5-year *p*-value |
| --- | --- | --- | --- |
| Image-only | 0.573 | 0.397 | 5.05E-13 |
| Clinical-only | 0.808 | 0.666 | 3.28E-14 |
| Image & clinical | 0.535 | 0.371 | 1.80E-22 |

**Table 4.** Survivorship of H/M/L patients of Wayne State data stratified by the TCGA-trained models.

| High/Moderate/Low | 3-year LR | 3-year HR | 5-year LR | 5-year HR | 3-year *p*-value | 5-year *p*-value |
| --- | --- | --- | --- | --- | --- | --- |
| Image-only | 0.858 | 0.857 | 0.662 | 0.646 | 5.00E-01 | 1.84E-01 |
| Clinical-only | 0.605 | 0.555 | 0.507 | 0.453 | 3.25E-03 | 2.14E-03 |
| Image & clinical | 0.747 | 0.605 | 0.682 | 0.545 | 1.14E-09 | 2.15E-05 |

**Fig. 5.**
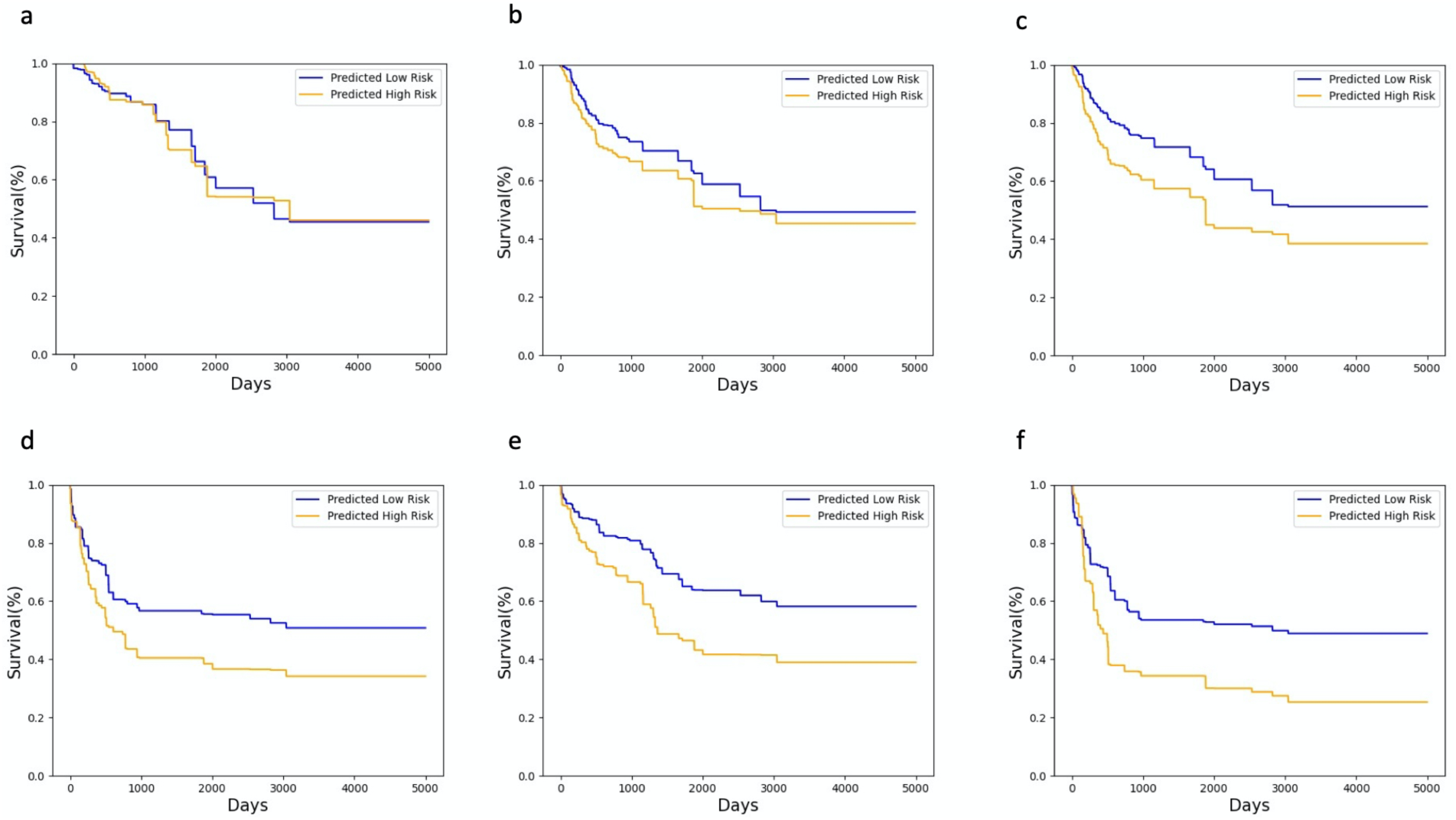
External validation using Wayne State data. a-c. The Kaplan-Meier curves of High/Moderate/Low patients using TCGA-trained classifiers for (a) image-only, (b) clinical-only, and (c) image & clinical models tested on external data. d-f. The Kaplan-Meier curves of High/Low patients using TCGA-trained classifiers for (d) image-only, (e) clinical-only, and (f) image & clinical models tested on external data.

We next considered the simpler problem in which only the HR and LR patients were in the test set. As expected we found better stratification than in the HR+LR+MR case. We observed significant stratifications for the image-only (3 year survival p-value=5.05e-13), clinical-only (3 year survival p-value=3.28e-14), and image & clinical models (3 year survival p-value=1.80e-22). Notably, the image & clinical model has performance superior to the image-only and clinical-only models (Figures 5d-f). Interestingly, although the image-only and image & clinical models performed similarly on TCGA test data (Table 1), the image & clinical model outperforms the image-only model on the Wayne State data (Table 3). This suggests the integrative model is more robust to stain differences across datasets.

Our pathologists further evaluated the heatmaps of the image & clinical model in the WSU cohort (see Supplementary Figure 2). These confirmed similar findings to the TCGA test set, i.e. that nuclear shape, nuclear size pleomorphism, intense cellularity, and abnormal structures are associated with high risk.

### 3.6 Robustness of separating moderate risk patients into high/low risk groups

We combined the Wayne State and TCGA data to more exhaustively investigate how MR patients can be computationally stratified into high and low risk groups. We used the image & clinical model and trained on HR and LR patients, analogous to Figure 2d. Given the small number of MR patients in the WSU cohort (N=9), we tested this in two ways: (1) training on WSU and testing on TCGA (N=45 in MR group), and (2) forming a combined multicenter dataset (TCGA+WSU) and testing/training on subsets.

First, we considered the model trained on WSU patients. We confirmed that the model trained on WSU HR and LR patients is able to effectively stratify a test set made of TCGA HR and LR patients (Supplementary Figure 6). We then tested how the model can stratify TCGA MR patients by risk. The model is able to stratify MR patients into higher risk and lower risk sets (5-year p-value= 0.03), though as expected stratification is not as distinct as for the HR/LR test sets. Second, we trained a model from the combined WSU+TCGA set. As expected, this model was able to stratify a reserved set of HR and LR patients by risk (Supplementary Figure 6). It also was able to separate MR patients into higher risk and lower risk (5-year p-value=5.63e-14), with a highly significant p-value. Interestingly, both of the MR stratification tests yielded long term survival ratio differences in the two predicted groups. Our results indicate that MR patients share enough similarities with HR and LR patients to improve survival stratification.

**Fig. 6.**
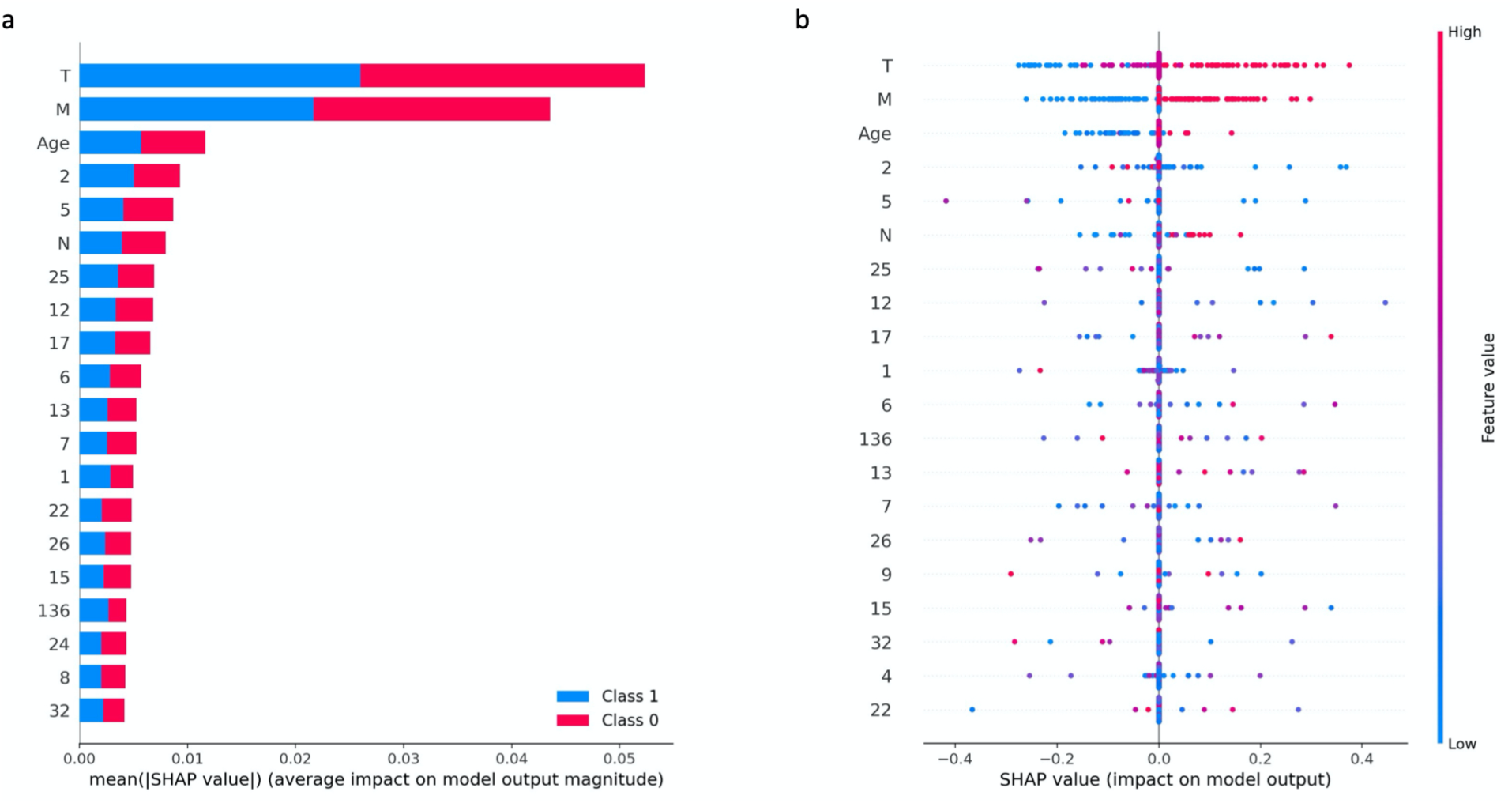
SHAP values of individual features of image & clinical model applied to Wayne State data. (a) Bar plot of the average SHAP values for top predicted features to illustrate global feature importance in class 1(High risk) and class 0 (Low risk). (b) SHAP values of top features across the Wayne State dataset. The plot sorts features by the sum of SHAP value magnitudes over all samples. The color represents the feature value (red high, blue low).

### 3.7 Feature importance for colon adenocarcinoma risk

To improve interpretability of our deep learning models, we used SHAP^44 45,46^ to measure the contribution of each clinical or Inception v3 image feature to the model output (see methods). We describe results for the model trained on TCGA and tested on Wayne State. We found that T stage, M stage, and age are the most impactful features in the integrative model (see methods, Figure 6). Although only two InceptionV3 features have comparable importance to these clinical variables, the total importance of InceptionV3 features (11.84) is higher than clinical variables (6.63). This may be explained by the fact that image contributions are spread across 2048 InceptionV3 features, while there are only 6 clinical variables for each patient. Interestingly, although individual clinical variables have high importance, the clinical-only model does not separate patients, suggesting the importance of cross-talk between clinical and image features.

## Discussion

### Integrative analysis improves stratification

While the utility of individual data modalities, such as clinical variables, mutation signatures, and WSIs, for patient stratification has been established^7-11,21,22,54,55^, our study demonstrates that integrative analysis improves patient risk stratification even for the challenging case of patients with intermediate survival times. Our image & clinical model showed more robustness to stain differences than the image-only model (section 3.5). Of potential importance is that cross-talk, i.e. variable-variable interactions, between image features and clinical variables, is informative of patient risk (see Tables 2 and 3; see Figures 2 and 5). Quantifying the crosstalk between each image feature and each clinical variable is an open research question for non-parametric deep learning models.

### Data heterogeneity requires flexible models and reliable training data

Our approach showed comparable performance even though we used a much smaller dataset size (231 patients) than other recent studies (>5000 patients^21^, >2800 patients^22^, >1000 patients^56^). We believe this is because of a number of strengths of our model. First, restricting image analysis to regions with high tumor content is crucial for reliable risk assessment (Results and Figure 4). Second, our training process uses classification-based losses and combines local tile-level with global patient-level information to improve model training. Classification based models are flexible, and are less prone to parameter estimation issues compared with those that directly use time-to-event values, e.g., the Cox proportional hazard model^57^. Third, restricting to patient subsets with strong survival differences produces a more reliable training dataset. While previous works have assigned a continuous risk score to all patients, e.g. Cox hazard ratio, and identified MR patients as a post-process^21^, our approach of binary classification of MR patients yields clearer results.

The combining of local tile-level image features with patient level information has theoretical advantages as it is a form of context-aware learning^32^. This approach is superior to^21^, where tile-level image features are first combined to a patient level image features, then patient level image and clinical variables are combined afterwards. Particularly, only our model was able to detect the importance of cross-talk between local image features and clinical variables. In ^21^, almost all of the signal was due to the image features (73-80%) with a lesser contribution from clinical features (T, N, and grade total: 18%) and no apparent cross-talk.

### Heatmaps enable automatic detection and interpretation of risk indicators

Prediction heatmaps from our computational model enable identifying regions that are informative of risk, improving model interpretability and discovery of novel prognostic markers. Specifically, our pathologist evaluations of the model predictions resulted in the findings that nuclear shape, nuclear size pleomorphism, intense cellularity, and abnormal structures are associated with higher risk (see Results and Figure 3). Our predictions also comport with known histopathological risk features. Histopathological tumor grading is used in the College of American Pathologists (CAP) protocol for colon cancer reporting as part of the diagnostic standard template, and has been shown to correlate with patient survival^49-52^. We observed a similar trend in both the TCGA-COAD and the WSU sets during annotation and clinical data collection. Such stratification based on histology is used in pathology reports as either a 2-tier, 3-tier, or 4-tier classification of tumors from well differentiated to poorly differentiated ^58^. Comparable patterns of high and low risk morphology were detected by the deep learning model as shown in Figures 3 and S2.

### Future directions

Our integrative deep learning model has potential to improve clinical decision making. For example, patients with a higher predicted risk of mortality may receive personalized treatment plans with closer follow ups. Patients considered under current standards to be moderate risk may especially benefit, as their outcomes are difficult to predict^59,60^ and better distinguishing their risk will be clinically useful^61 62^. Computational risk prediction may also present evidence for improved approval of expensive scans and improved patient counseling. To realize these translational goals, an important future direction will be to further explore the cross-talk between morphological features and clinical variables. Recent studies suggest that individual image deep learning features encode interpretable morphologies^63^ and that small clusters of deep learning features encode distinct markers of risk^21^. Cross-talk can be further studied by identifying the distinct morphologies encoded by deep learning features in regions of interest, and then evaluating correlations between deep learning features and clinical variables in each risk group. Finally, while the current study establishes the utility of integrative models in stratifying moderate risk patients, given the small portion of such patients in our datasets, larger datasets are necessary to improve risk predictions.

## Data/Code Availability

WSU Data and code are available upon request. TCGA data is publicly available and can be downloaded from the GDC portal (https://portal.gdc.cancer.gov/).

## Acknowledgements

JHC acknowledges support from NCI grant R01CA230031 and P30CA034196. AF acknowledges support from a JAX Scholar award.

