## Supplementary figures for "Integrative deep learning analysis improves colon adenocarcinoma patient stratification at risk for mortality": Supplementary.docx

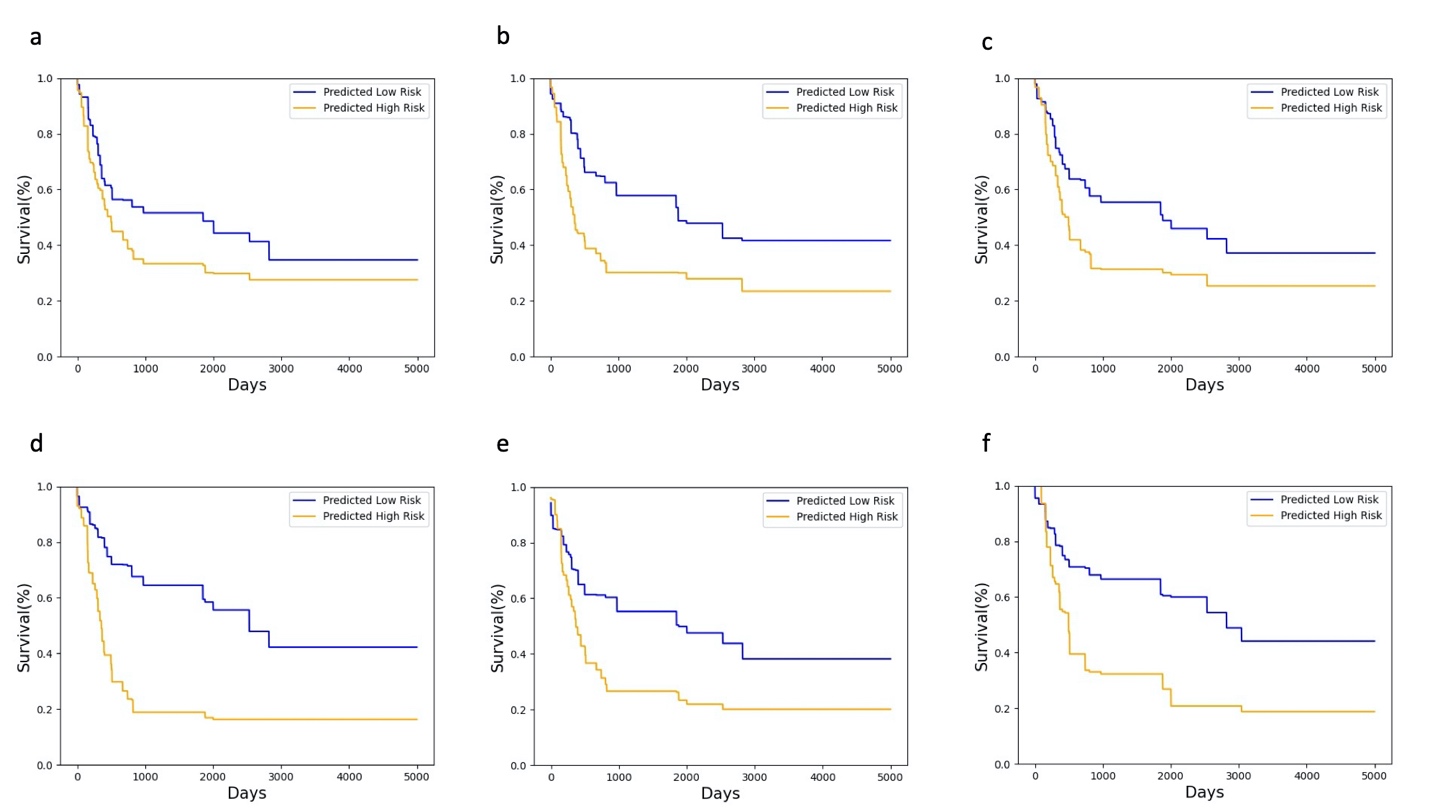

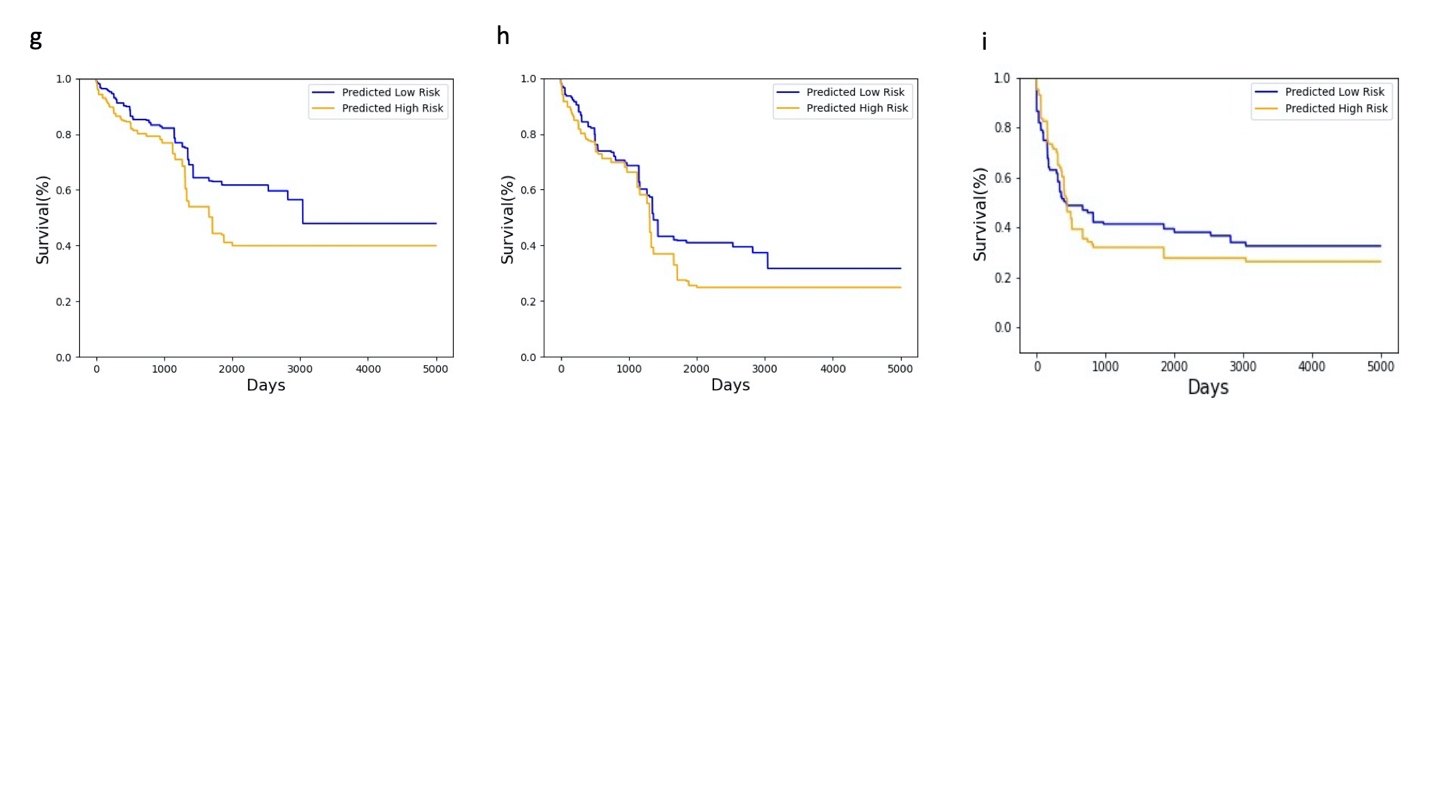

**Fig. S1 WSIs are informative of risk.** K-M curves of H/L groups using the (a) mutation, (b) clinical, (c) clinical & mutation, (d) image & clinical, (e) image & mutation, (f) image & clinical & mutation models. K-M plots of H/M/L patients for (g) clinical, and (h) clinical & mutation models. (i) K-M plots of H/L patients for deep learning Cox model using image features.

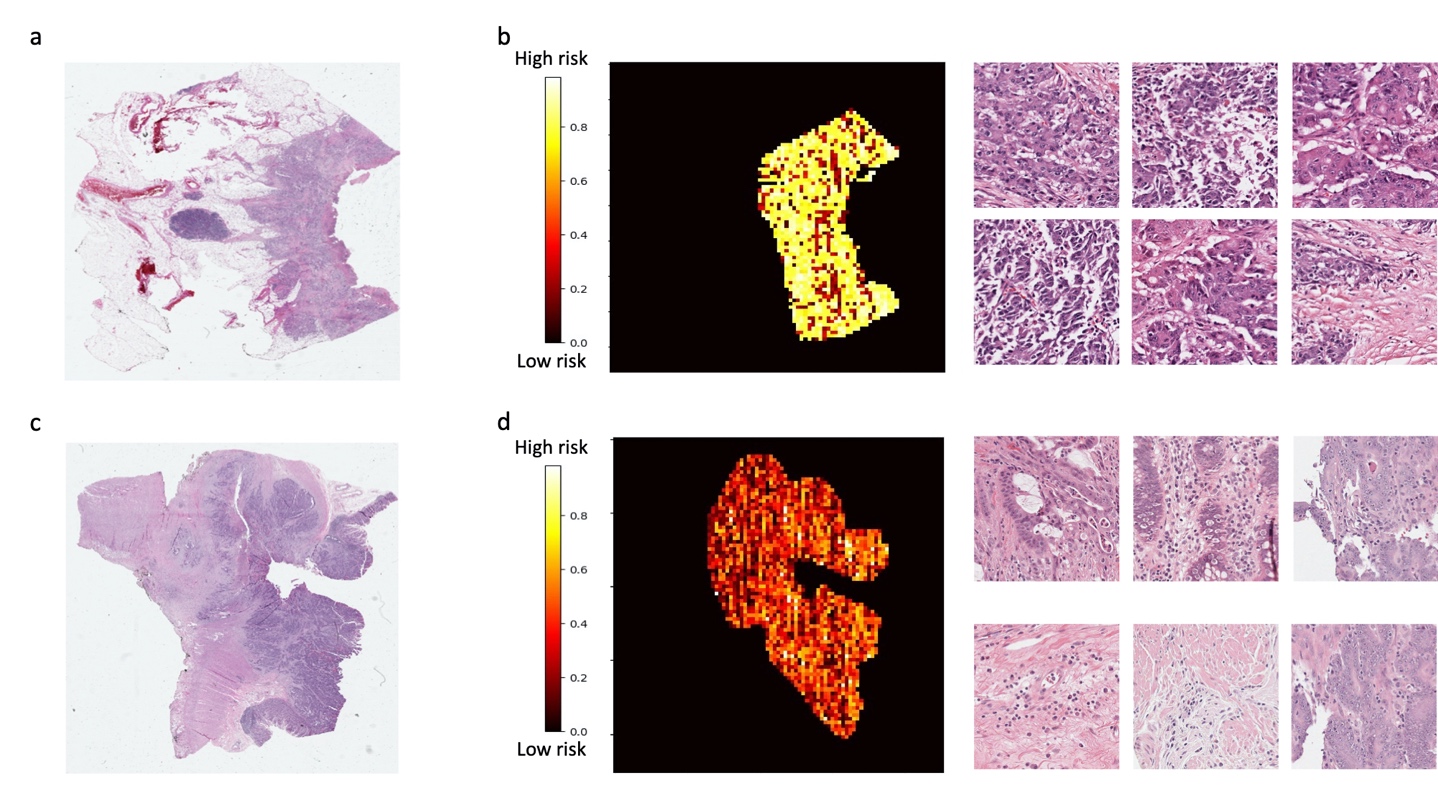

**Fig. S2 Representative H&E slides from Wayne State data and their predicted heatmaps.** WSIs of a (a) high risk patient and (c) a low risk patient. The prediction heatmaps of a (b-left) high risk patient and (d-left) low risk patient. Example tiles predicted as (b-right) high risk and (d-right) low risk.

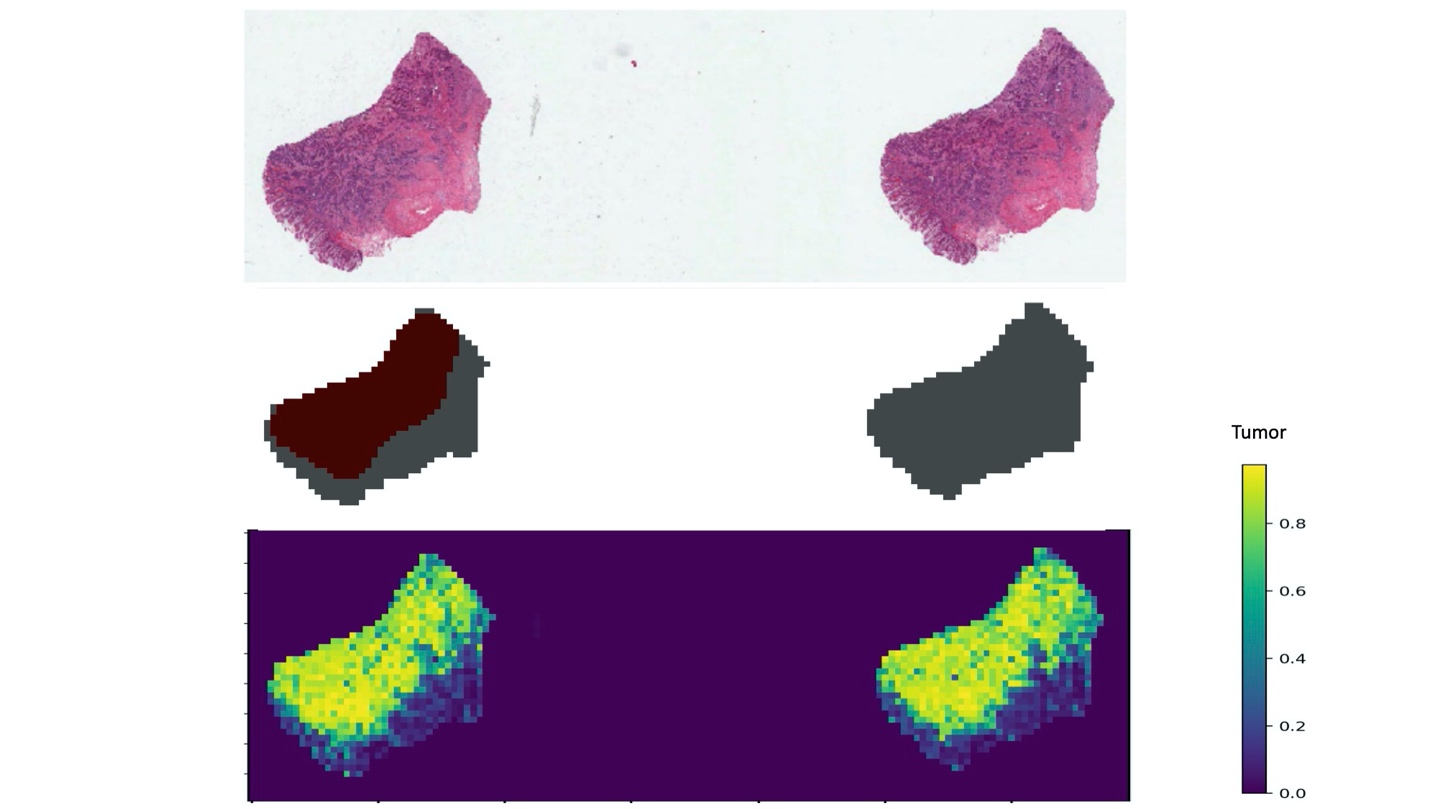

**Fig. S3 Computational tumor detector identifies unannotated tumor regions.** WSI of a TCGA patient. Left and right pieces are from adjacent slices of the same tumor. (top). (Middle) Red regions show the tumor annotations generated from pathologists. (Bottom) Tumor prediction heatmaps. Yellowish tiles outside the pathologist annotations tend to be regions with low but non-zero tumor content. Also, the right slice was not selected for manual annotation but the detector correctly predicts a similar tumor localization as in the adjacent slice.

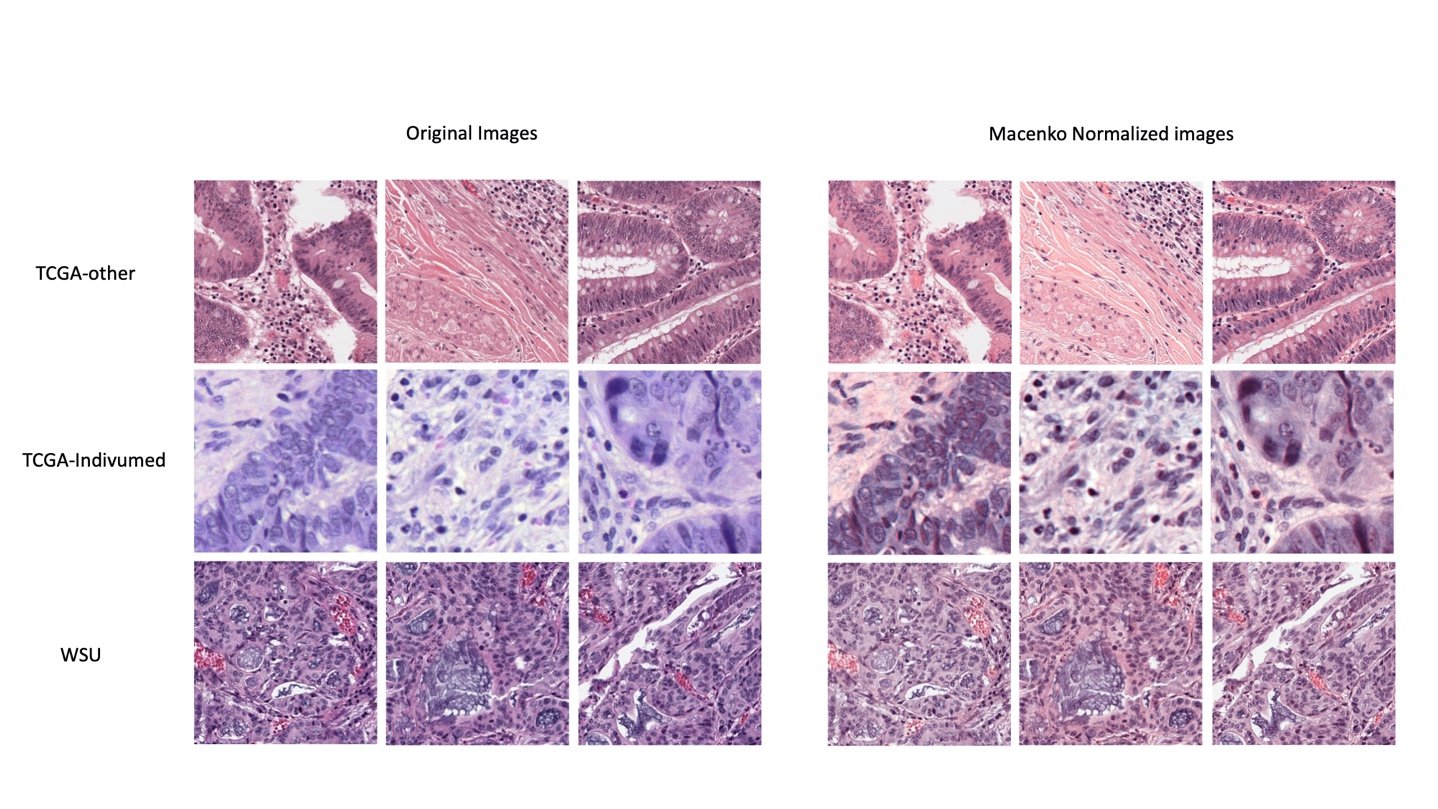

**Fig. S4 Macenko normalization of COAD tumor tiles from multi-center WSIs.** Left panel shows the original COAD images from TCGA Indivumed institute (top), WSU(middle) and TCGA-other institute(bottom). Right panel shows the Macenko normalized COAD images from TCGA Indivumed institute (top), WSU(middle) and TCGA-other institute(bottom)

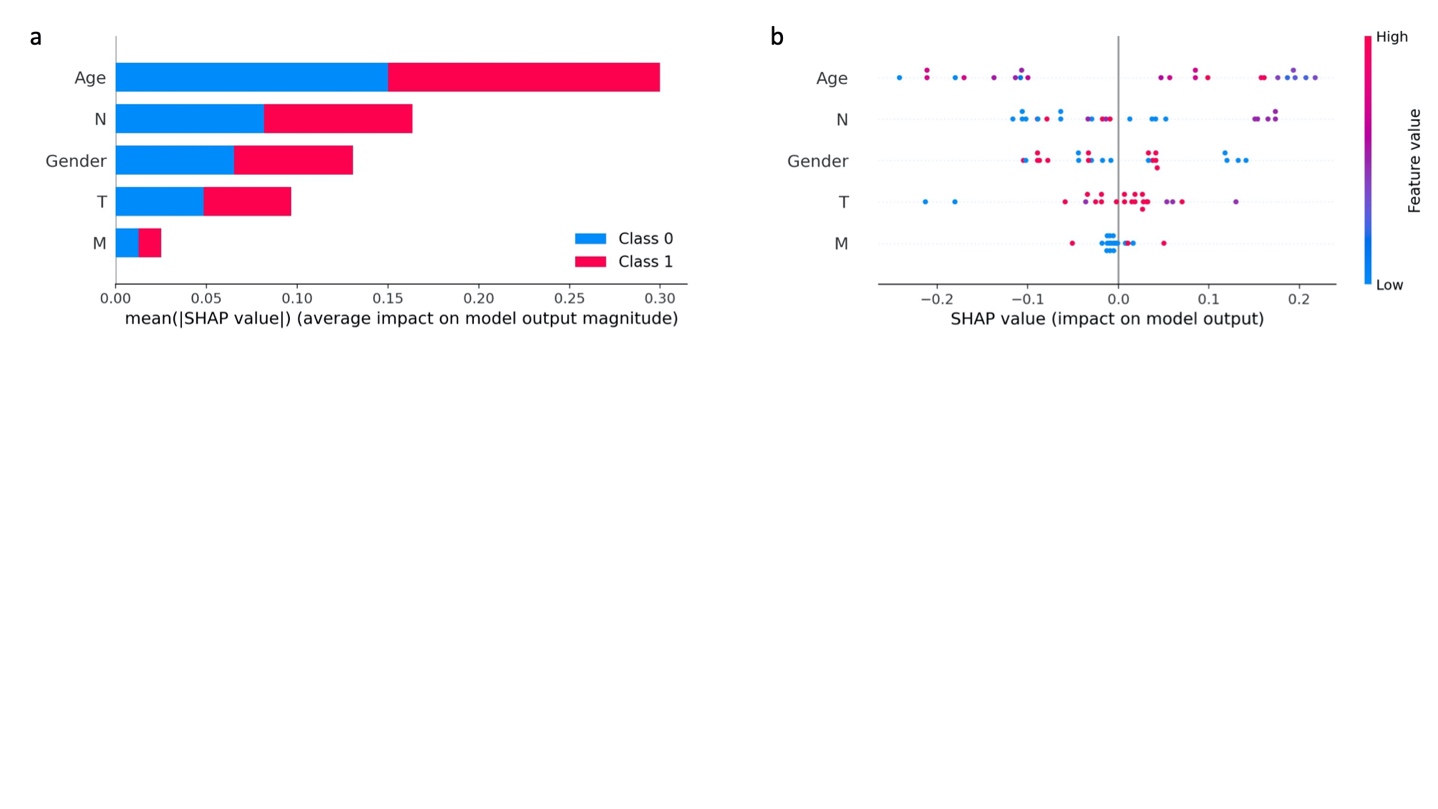

**Fig. S5 SHAP values of individual features of the clinical-only model applied to Wayne State data. (**a) Bar plot of the absolute SHAP mean values for top predicted features to illustrate global feature importance in class 1(High risk) and class 0 (Low risk). (b) SHAP values of every feature for every sample. The plot sorts features by the sum of SHAP value magnitudes over all samples. The color represents the feature value (red high, blue low).

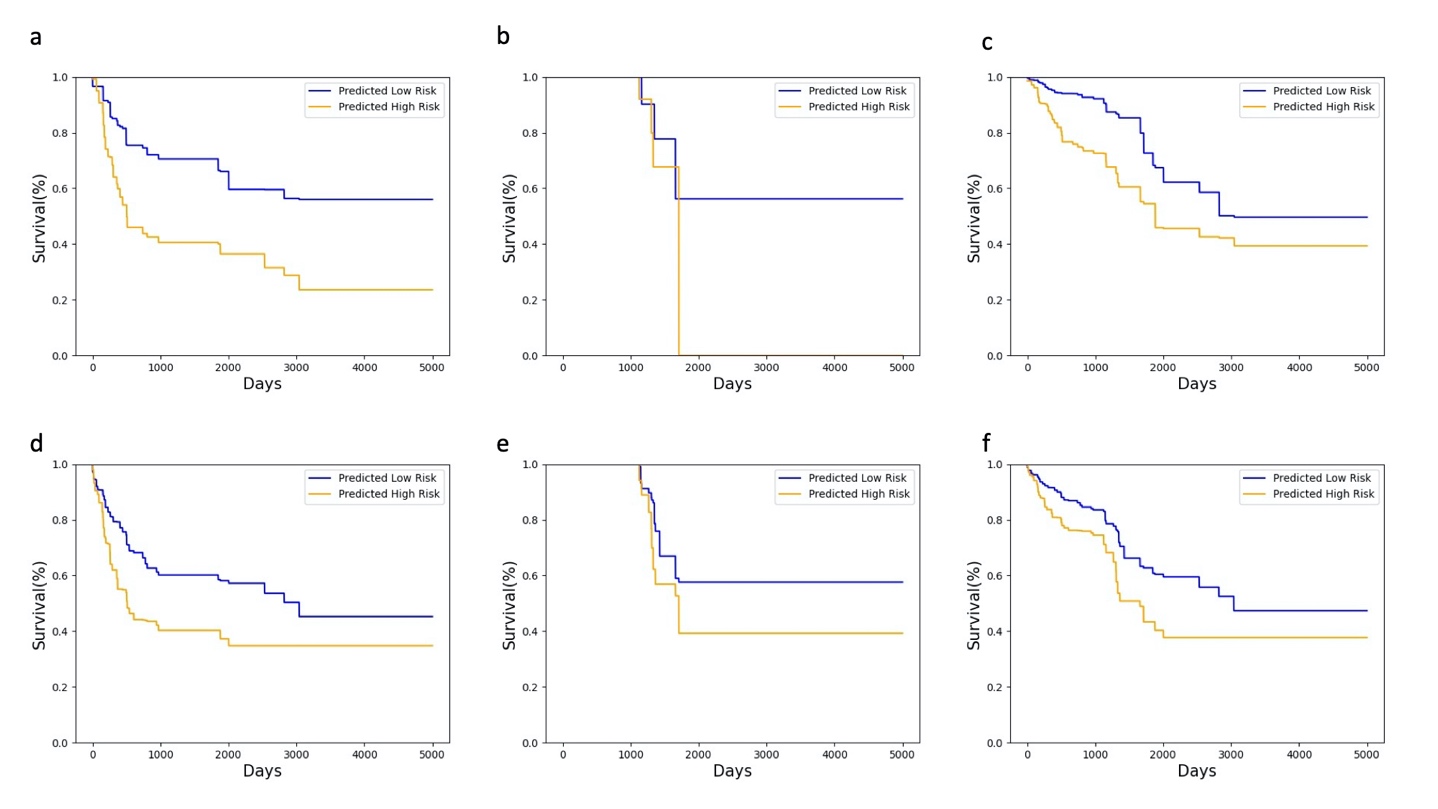

**Fig. S6 Robustness of splitting of moderate risk patients into higher or lower risk groups.** a-c. The K-M curves of classifiers trained on WSU and tested on TCGA for (a) High/Low patients, (b) Moderate patients, (c) High/Moderate/Low patients. d-f. The K-M curves of classifiers trained on TCGA+WSU and tested on TCGA+WSU for (d) High/Low patients, (e) Moderate patients, (f) High/Moderate/Low patients.

**Supplementary Tables**

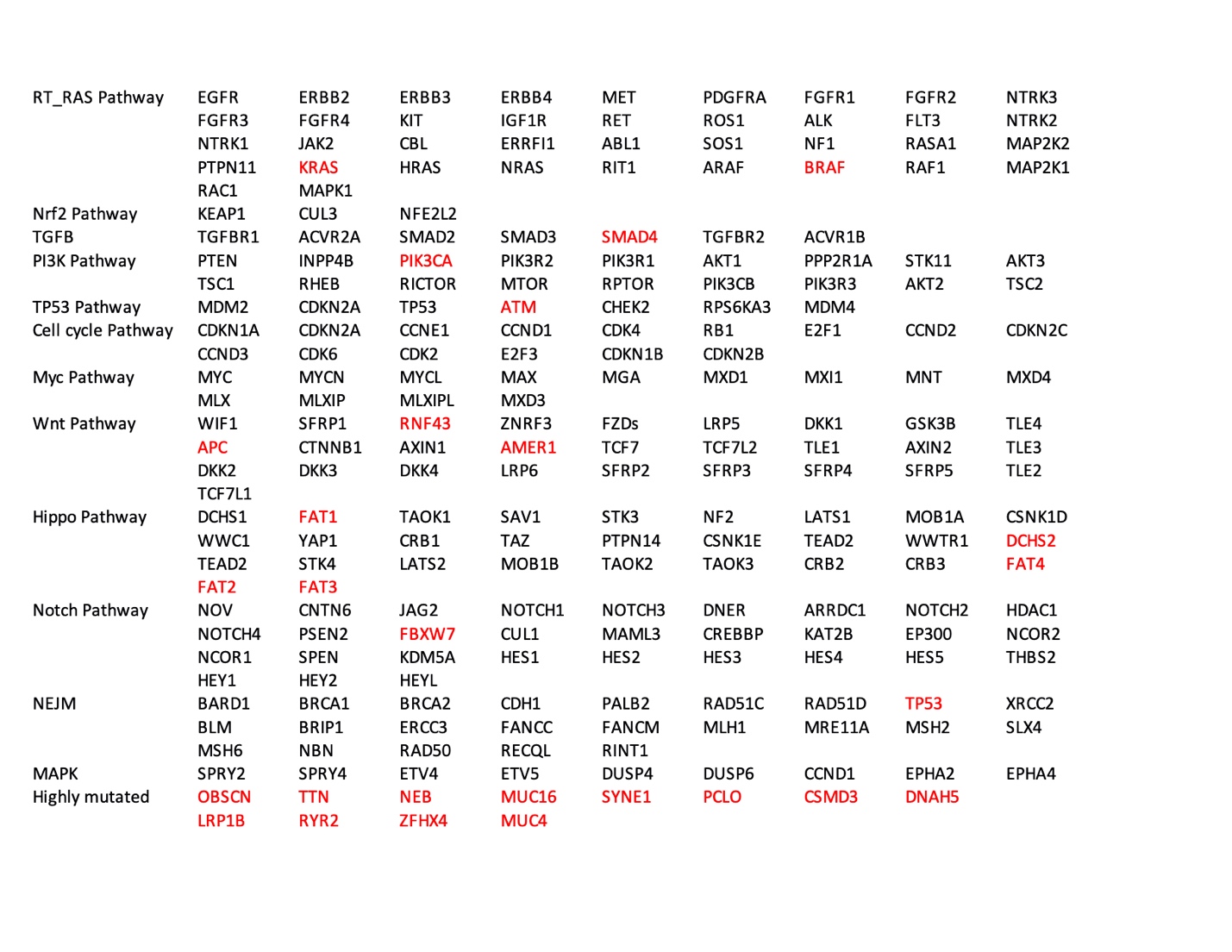

**Table. S1 Gene list**. Candidate genes based on literature review. Only genes with mutation rate >10% (highlighted in red) were used.
